# <u>10 Simple Rules</u> for Sharing Human Genomic Data

**DOI:** 10.1101/094110

**Authors:** Manuel Corpas, Charlotte Whicher, Nadezda V. Kovalevskaya, Tom Byers, Amanda A. McMurray, Fiona G.G. Nielsen, Varsha K. Khodiyar

## Introduction

Delivery of the promise of precision medicine relies heavily on human genomic data sharing. Sharing genome data generated through publicly funded projects maximises return on investment from taxpayer funds and increases the likelihood of obtaining funding in future rounds [1]. More importantly, genome data sharing makes it possible for other scientists to reuse existing datasets for further research and constitutes a direct measure of the current advancement in risk prediction, diagnosis, and treatment for genomic disorders [2].

Sharing of human genomic data carries responsibilities to protect confidentiality and the privacy of research participants [3]. In certain cases data sharing may be complicated or limited by agreements with outside collaborators or Institutional Review Board (IRB) rules. Despite this, there is a broad agreement among funders (e.g., NIH [4], Wellcome Trust [5] and many others) for investigators to share data broadly for secondary research purposes, in all cases consistent with applicable laws, regulations and policies. Human genomic datasets may be accompanied by sensitive clinical metadata from patients, including pictures, medical history, sex, age, etc., which may have a critical role in diagnostics and generation of actionable outcomes.

Usually, disease causing mutations on their own are not a threat to the privacy of patients. However, it was shown by Homer et al [6] that making human genome data publicly available in an anonymised form does not completely conceal identity, since it is straightforward to assess the probability that a person or relative participated in a study, particularly when phenotype and clinical metadata are also available. A large number of studies have suggested that a technically sophisticated data breach may exploit a wide range of human genetic data [2]. Therefore, enabling ethical data sharing while ensuring genetic privacy remains challenging.

There are a number of international efforts underway to establish standards for good practice in sharing human genomic data. The FAIR (Findable, Accessible, Interoperable and Reusable) principles provide a general framework for data sharing that are also applicable to biomedical research data [7]. The Global Alliance for Genomics and Health has a clinical working group to develop compatible, readily accessible, and scalable approaches for sharing clinical data and linking it with genomic data [8]. Specialised data repositories are also being developed to facilitate the sharing of clinical data that cannot be openly shared for reasons of patient privacy [9]. Specialised data journals such as *Scientific Data* [10], *GigaScience* [11] and *Human Genome Variation* [12] enforce best practice for publishing data, whilst providing an incentive for researchers to share their data via a data paper.

These 10 Simple Rules have been developed from our combined experiences of working with human genomic data, data repositories and data users. We do not claim that these rules will eliminate every possible risk of data misuse. Rather, we hope that these will help researchers to increase the reusability of their human genomic data, whilst also ensuring that the privacy of their subjects is maintained according to their consent frameworks. Many of the principles presented are also applicable to other types of clinical research data, where participant privacy is a concern.

### Rule 1: Recognise the intrinsic value of the data

Genomic data has great value. An individual’s genomic data has value to that individual and their biological relatives. An individual’s genomic data also has value to the research community, as a unique dataset and as part of a larger dataset composed of multiple individual genome sequences. In the UK, there is currently a moratorium on the use of individual’s genetic data for life insurance purposes [13], but as a responsible researcher, it is important to recognise that an individual’s genomic data can have commercial value too. A number of companies are commercialising human genome datasets. For instance, 23andMe sells access to the database created by customers who have bought 23andMe’s DNA test kits and donated their genetic and health data for research [14]. Seven Bridges provides “immediate access to some of the world’s richest [genomics] datasets” [15].

Making research data available for reuse is an important component of reproducible research [16]; however, the intrinsic value of genomics data to multiple parties, requires that sharing these types of data merits specific consideration.

### Rule 2: Choose the appropriate patient consent framework

Obtaining appropriate consent to collect genotype, phenotype and any other type of human data is usually the responsibility of the researcher or clinician who carries out the study. Consent forms should specify the goals of the immediate project, and explicitly describe in clear terms if the data are intended to be shared beyond the current scope of the project. If wider data sharing is intended, the consent form should clearly state potential risks and benefits to the participants, as well as any data anonymisation procedures that will be undertaken.

Different levels of anonymisation are possible and the suitability of these should be considered prior to data collection. Normally the approval of an Institutional Review Board (IRB) should be sought prior to data collection. Different consent forms provide varying degrees of identity exposure from the individual. For example, the Personal Genomes Project, provides complete access of identity and traits from the study participants [17] under a CC0 license waiver [18]. This consent framework is not, however, the usual one when dealing with human genome data derived from the clinic. NIH funded studies require third party researchers to describe how they intend to use the data through a Data Access Request and, through a Data Use Certification Agreement, are requested to adhere to the NIH Genomic Data Sharing Policy’s ethical principles, terms of data access, and privacy safeguards [19]. In the UK, Genomics England consent forms are classified according to patients being affected with cancer or rare diseases and the consent framework allows access to summary statistics in a controlled environment [20].

### Rule 3: Check whether support for data sharing is available

There are resources [21] that list the major funders research policies, such as NIH, Wellcome Trust and ERC. Funders like these can provide services to help scientists to design a data sharing plan and to decide where to share datasets. Some may also offer the services of a data scientist to support the data sharing process and, in some circumstances, they may be able to provide additional funds to cover data deposition to an appropriate repository.

### Rule 4: Understand the datasets you generate

There are two key properties to consider prior to data generation, the size of the data set and the format in which the data should be shared. The size of the final dataset may impact on the type of repository that can be used to share the data (see Rule 8). Considering the most appropriate format for both the raw and processed data, will impact on the ease with which the data can be shared.

### Rule 5: Context is king: adhere to metadata standard descriptions

Metadata determines how visible a dataset will be, not only to users searching for it but also to search engines and crawling bots. Use of the most appropriate standard for metadata descriptions will increase its visibility and reusability [22]. For example the use of MIAME (Minimum Information About a Microarray Experiment) for microarray data [23], increases the discoverability of microarray data in MIAME-compliant repositories such as NCBI GEO [24] and EBI ArrayExpress [25].

### Rule 6: Check the accuracy of your metadata

Include relevant keywords in dataset metadata descriptions. For example if healthy individuals are involved, ensure that the keyword “healthy” is included. If disease phenotypes are associated with genome data, consider annotating variant files with keywords such as those provided by the Human Phenotype Ontology [26]. This will help find your data when searching for specific genetic conditions. For datasets that include different data types, the metadata should include the file types describing those data types (e.g. VCF [27], SRA [24], BAM/SAM [28], FastQ, etc.).

### Rule 7: Maximise the machine readability of your metadata

Maximising the likelihood that data can be discovered is a vital component of the data sharing process. Writing a data paper will increase the discoverability of the dataset, since the article will be indexed in bibliographic databases such as PubMed. It is also worthwhile to plan for ways in which the data itself can be made discoverable, as well as considering both the human and machine accessibility of the dataset. For example, it was recently shown that human gene symbols were converted to dates in the supplementary data files of some published papers [29], which meant these gene symbols were not machine readable. Simple strategies can avoid such errors by including data units in tables and keeping data types consistent across columns or rows to avoid mixing of strings with numbers. Avoid the use of acronyms where possible, and make sure they are defined if their use is unavoidable.

### Rule 8: Choose the most appropriate repository for your data

Using specialist data repositories for research data, helps to ensure that this data is archived and preserved in a data type-specific way. For instance, array-based human data would usually be submitted to repositories such as GEO [24] or ArrayExpress [25], while unaligned raw sequence data should usually go to repositories such as SRA [24] or ENA [30]. For clinical genomics dataset deposition, the European Genome-phenome Archive (EGA) and the NCBI equivalent dbGaP provide controlled access data storage. Both resources allow submission of sequence, array-based and phenotypes.

However, as mentioned in Rule 1, human genomic data merits specific consideration of participant privacy. Care must be taken to archive such data in repositories that have workflows in place to ensure data access is only given to those that fulfill the relevant requirements. There are a number of repositories which are suitable for any human genomic data [9], and there are also specialist repository indexing services which can help to guide an informed choice [22].

### Rule 9: Upload both raw and processed data

Both raw and processed data are valuable resources. However, processed data should never be assumed to be sufficient for reanalysis of specific results. Some users want processed data because they do not have the resources to extract it from the raw data. However others will prefer to process the raw data into their own pipelines and adjust it to their chosen parameters or thresholds. Thus when working with NGS human data, it would be advisable to include all fastQ, BAM and VCF files, together with all the metadata needed in order to reproduce the same processing that lead to the VCF files.

### Rule 10: Make it easy to cite your data

Data citation is growing in importance as a way in which researchers can gain recognition for making data available, as well as providing provenance for the data. Therefore it is important to ensure that data is shared in a citable way. A good data repository will provide a persistent and unique identifier for each data archive, enabling data to be cited. The main NCBI and EBI databases use accession identifiers, and other repositories may use DataCite digital object identifiers (DOIs). Both accession IDs and DOIs can be cited in scholarly works as, for example, can be seen in the guidance *Scientific Data* provides to its authors on how to cite data [31].

## Conclusion

We have presented the 10 Simple Rules that we recommend to maintain best practice when sharing human genomic data. As personalised medicine starts to impact patients, it is expected that datasets containing potentially sensitive information will become more widespread. Hence having a set of guiding rules that help keep patient data reusable whilst complying with patient consent around sharing of their data is crucial, if we are to leverage the power of NGS data from human origin and so realise the promise of precision medicine.

## Acknowledgements

We are grateful to the public repositories that make it possible for scientists to upload their genomic datasets at no cost.

